# Comparative analysis of protease-inhibitor gene family counts under different counting units and genome annotations

**DOI:** 10.64898/2026.09.22.753331

**Authors:** Fang Zhao, Juan Zhao, Feng Zhao

## Abstract

**Background:** Gene family sizes are often compared across species using the protein sets released with genome assemblies. However, counts of domain-positive protein entries can differ from counts of genes when an annotation contains several entries per gene, and genomes are annotated by different pipelines. In this study, we measured how these two factors change the counts of protease-inhibitor domain families.

**Results:** Seven protease-inhibitor families were counted in eight animal genomes. Collapsing protein entries to genes reduced the counts in 36 of 48 species–family combinations, by up to 4.6-fold (Kunitz/BPTI in *Aedes aegypti*, 23 entries in 5 genes), and reversed the order of two species in 9 of 196 pairwise comparisons. In 218 of 223 gene–family records with several positive entries, all entries shared CDS coordinates with the representative entry. Two annotations of the same leech (*Hirudinaria manillensis*) assembly differed in a family-specific way. In the published annotation, 38 of 50 Antistasin-positive genes and both Hirudin-positive genes had no same-strand gene model in a second annotation, whereas most Kunitz/BPTI, Serpin, TIL and Thyroglobulin genes had one-to-one partners. A search of the eight assemblies without gene models detected all 338 annotated Kunitz/BPTI-positive genes when the queries included proteins of the same species. Queries from other species recovered only 31–83% of these genes, and Kunitz-like candidate regions outside the annotations were most numerous in the tick *Ixodes scapularis*. For the mosquito and the tick, the second public annotations shared nearly all gene-model coordinates with RefSeq and were therefore excluded from the comparison of annotation sources.

**Conclusions:** These results indicate that gene family counts depend on the counting unit and on the annotation source. Comparisons across species should count each gene once and state how domain positivity was assigned to genes. In addition, the source and version of each annotation should be reported, the annotations should be checked against the genome assembly, and the overlap of gene models should be examined before two annotations are compared.

## Background

Gene family sizes are commonly compared between genomes. In leeches, for example, increases in the copy number of anticoagulation-related genes have been linked to blood feeding [1]. Such sizes are usually derived from the protein sets released with genome assemblies [1, 2].

Two features of public annotations can change such counts. First, a protein set usually contains one entry for each annotated transcript, so counts of domain-positive protein entries can differ from counts of genes. The number of protein entries per gene differs between annotation projects, and in the eight genomes analysed here it ranged from 1.00 to 2.67 (Table 1). For orthology inference, one representative transcript per gene, usually the longest, is commonly used [3]. Second, genomes are annotated by different pipelines with different evidence, so a family that is well captured in one genome may be partly missed in another [4, 5]. Small families of short, divergent secreted proteins are expected to be most sensitive to both effects, because a few extra transcript models or a few missed genes change the count substantially.

**Table 1.**
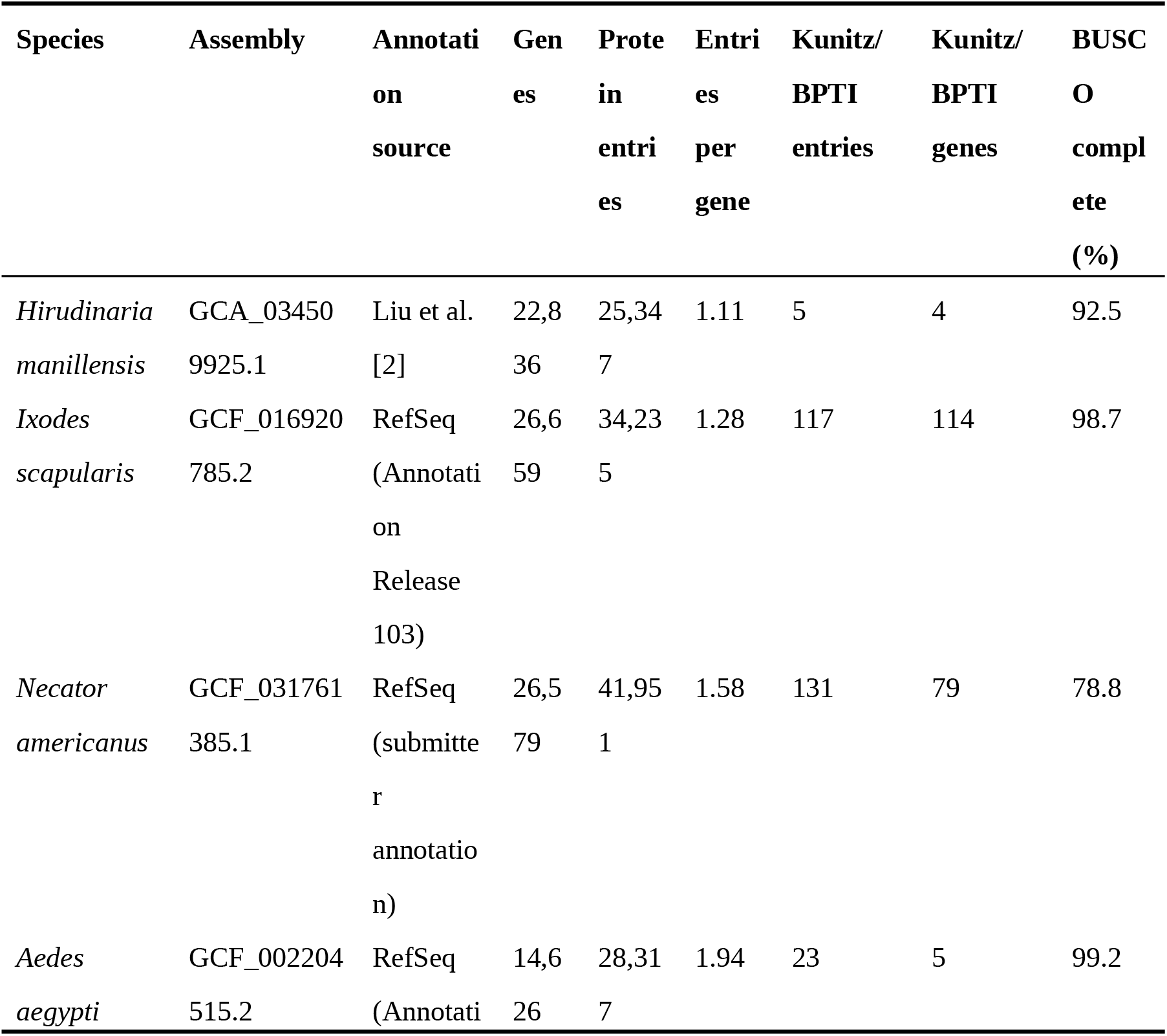
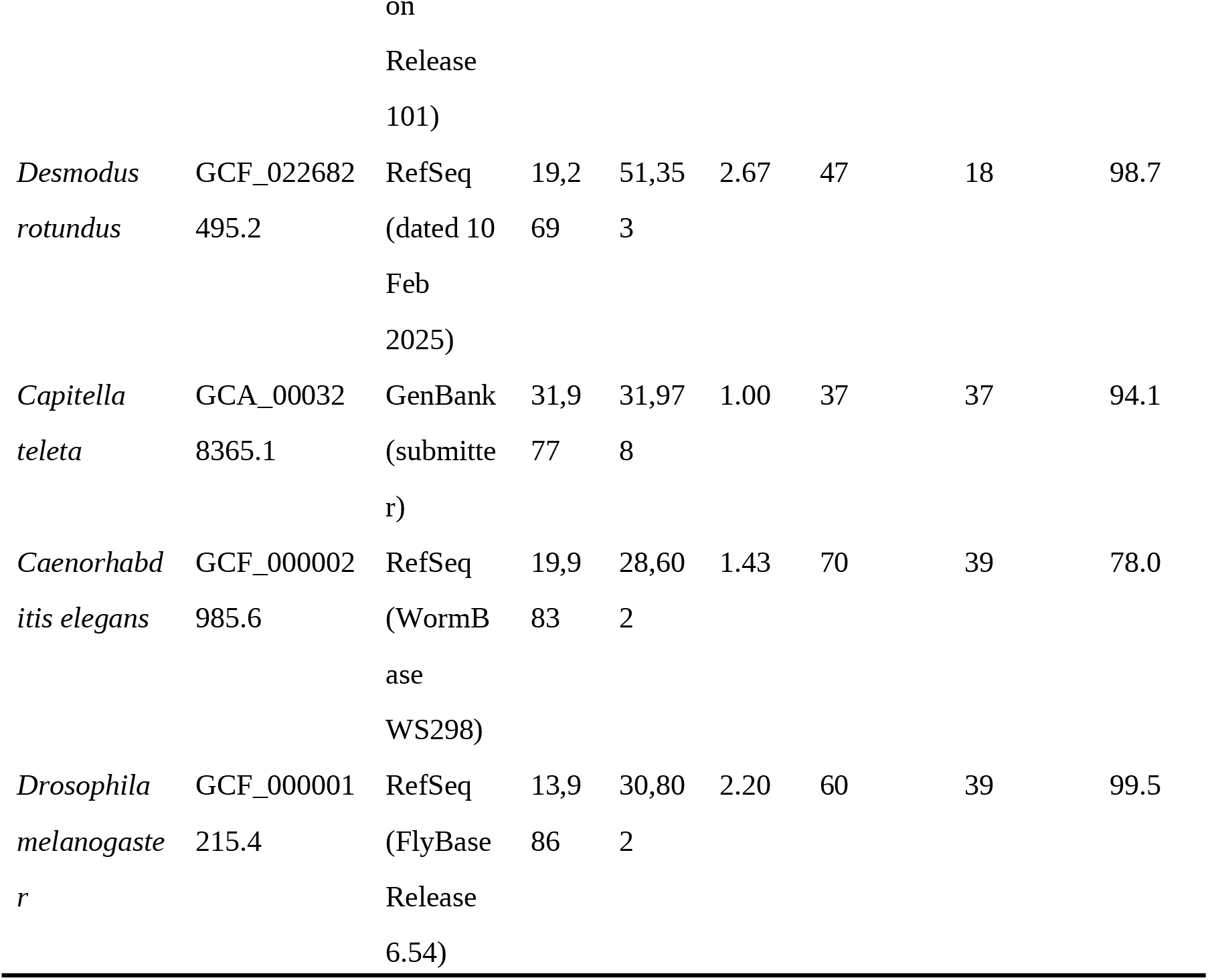
Genomes, annotations and Kunitz/BPTI counts. Genes and protein entries refer to the full annotated protein set. Kunitz/BPTI genes are genes whose longest entry carries the domain. Annotations were downloaded on 16 July 2026. BUSCO 6.1.0 completeness (metazoa_odb10, protein mode) was assessed on the longest entry of each gene.

Protease-inhibitor domain families are a useful case for measuring these effects. Families such as Kunitz/BPTI, Kazal, Antistasin and Hirudin are small and often divergent, and many of their members have been described in salivary transcriptomes and genomes of blood-feeding animals [6, 7]. The Kunitz/BPTI family has expanded in the tick *Ixodes scapularis*, and many of its members are expressed in the salivary glands [8]. Protease inhibitors of several families have also been described in salivary transcriptomes of mosquitoes and the vampire bat [7, 9]. The genomes of these animals have been annotated by RefSeq, by the authors of individual genome projects and by other pipelines. For the medicinal leech *Hirudinaria manillensis*, two genome projects have reported counts of antithrombotic genes [2, 10]. In addition to the published annotation of one of these assemblies [2], a second annotation of the same assembly was generated in this study.

In this study, we used seven Pfam domain families associated with protease inhibitors as a test case to measure how the counting unit and the annotation source change gene family counts. The seven families were Kunitz/BPTI, Antistasin, Hirudin, Kazal, Serpin, Thyroglobulin type-1 and TIL. The eight genomes were chosen because their annotations come from different sources. We compared three counting units in the eight genomes, separated annotated transcripts from neighbouring duplicate genes, and compared two annotations of the same leech assembly. We also searched the genome assemblies for Kunitz/BPTI domains outside the annotations and examined the second public annotations of the mosquito and tick assemblies.

## Methods

### Genomes, annotations and protein sets

Eight annotated animal genomes were analysed: the leech *Hirudinaria manillensis* (GCA_034509925.1), the tick *Ixodes scapularis* (GCF_016920785.2), the hookworm *Necator americanus* (GCF_031761385.1), the mosquito *Aedes aegypti* (GCF_002204515.2), the vampire bat *Desmodus rotundus* (GCF_022682495.2), the polychaete *Capitella teleta* (GCA_000328365.1), the nematode *Caenorhabditis elegans* (GCF_000002985.6) and the fruit fly *Drosophila melanogaster* (GCF_000001215.4). Protein sets and GFF3 annotations of the other seven species were downloaded from NCBI on 16 July 2026. Six are RefSeq annotations [11]: NCBI Annotation Releases 103 (*I. scapularis*) and 101 (*A. aegypti*), the NCBI annotation dated 10 February 2025 (*D. rotundus*), and the submitter, WormBase WS298 and FlyBase Release 6.54 annotations distributed through RefSeq (*N. americanus*, *C. elegans* and *D. melanogaster*, respectively). The *C. teleta* annotation is the GenBank annotation of the submitters (Table 1).

No annotation is distributed with the leech assembly at NCBI. For this species, the coding sequences released by Liu et al. [2] were translated with the standard genetic code, and the GFF3 file of the same release was used for coordinates. This annotation combines a BRAKER track (22,764 gene models) with a second track of 72 manually curated antithrombotic gene models. A second annotation of the same leech assembly was generated in this study. Training genes and hints were derived with GALBA 1.0.11 [12] from 55,346 proteins of two other annelids, *Helobdella robusta* (RefSeq GCF_000326865.2, 23,368 sequences) and *C. teleta* (31,978 sequences). No *H. manillensis* sequence was used. AUGUSTUS 3.5.0 was trained with 942 complete gene models, with the stop codon excluded from the CDS (stopCodonExcludedFromCDS=true), and was run on each assembly sequence with the GALBA hints and extrinsic configuration. The merged prediction contained 20,377 genes and 21,805 transcripts (Additional file 1).

### Domain detection and counting units

Seven Pfam [13] profile HMMs were used: Kunitz_BPTI (PF00014.30), Antistasin (PF02822.20), Hirudin (PF00713.23), Kazal_1 (PF00050.27), Serpin (PF00079.26), Thyroglobulin_1 (PF00086.24) and TIL (PF01826.24). Domains were detected with hmmsearch (HMMER 3.4 [14]) using the family gathering thresholds (--cut_ga). Each protein entry was assigned to a gene from the CDS features of the GFF3 file, using the locus_tag, then the GeneID cross-reference and then the gene symbol; transcript identifiers were ignored. For the leech, the gene identifier given in each protein header was used. All protein entries were assigned to a gene in all eight species. Completeness of each protein set was assessed with BUSCO 6.1.0 [15] in protein mode with the metazoa_odb10 dataset (954 BUSCOs), using the longest entry of each gene.

For each family and species, three counts were recorded: (1) the number of protein entries carrying the domain; (2) the number of genes with at least one domain-positive entry; and (3) the number of genes whose longest entry carries the domain. The longest entry of each gene was used as its representative; when two entries had the same length, the first one in the file was kept. For every pair of species, the order of the two species under count (1) was compared with the order under count (3).

### Protein entries and neighbouring duplicates

For genes with two or more domain-positive entries, the CDS coordinates of each entry were compared with those of the representative entry of the same gene. Entries with identical amino acid sequences were also counted. To identify recent duplicates among different genes, the representative proteins of each family were compared with BLASTP 2.17.1+ [16] (E-value ≤ 1e-5, one high-scoring segment pair per protein pair). A pair of genes was recorded as a candidate tandem duplicate when the two genes were on the same sequence, were non-overlapping and were separated by no more than 20 kb. Their proteins also had to share at least 98% identity over at least 90% of the shorter protein. A second, more permissive criterion (90% identity, 50 kb) was used as a sensitivity analysis. Overlapping gene models with similar proteins were recorded separately, and the gene counts were left unchanged.

### Comparison of two annotations of the same assembly

The two leech annotations were compared by genomic position. The two annotations use the same 23 assembly sequences under different names, which were matched by identical sequence content (Additional file 1). For each gene, the span of all its CDS features was used as its locus. Genes were scored as domain-positive when any of their entries carried the domain. Within each family, positive genes from the two annotations were linked when their loci overlapped by at least 1 bp on the same strand. The resulting connected groups were classified as one-to-one, one-to-many, many-to-one or many-to-many. Positive genes without a positive partner were further classified by whether the other annotation had a gene model on the same strand at that position. These classes describe only the overlap of CDS spans, and a structural comparison would be required to show whether gene models were split, merged or truncated. Positive genes of the curated antithrombotic track that overlapped a positive gene of the BRAKER track were also recorded.

### Genome search for Kunitz/BPTI candidate regions outside the annotations

To find Kunitz/BPTI loci missing from the annotations, the eight genome assemblies were searched without using their gene models. The query set contained all Kunitz/BPTI-positive protein entries of the eight species, with identical sequences removed. Queries were aligned to each assembly with miniprot 0.18 [17] (-N 50 --outs=0.5), and the translated alignments were scanned with the Kunitz_BPTI profile HMM at the gathering threshold. The amino acid positions of each domain were converted to genomic coordinates with the alignment CIGAR string, so that frameshifts and introns were taken into account, and each domain was kept as a set of genomic segments. In addition, each assembly was translated in six frames with esl-translate (open reading frames of at least 20 amino acids) and scanned with the same profile. Hits with a sequence E-value and a conditional domain E-value of 1e-3 or lower were also recorded; hits passing either threshold were combined.

In this analysis, a gene was treated as Kunitz/BPTI-positive when any of its entries carried the domain. An annotated Kunitz/BPTI-positive gene was counted as detected when a domain segment overlapped one of its CDS segments on the same strand. The remaining domain hits were grouped by sequence and strand and merged when their segments overlapped. Each resulting region was assigned to one of three classes: regions on the same-strand CDS of a gene model without the domain; regions within the CDS span of a gene model on either strand but not on its same-strand CDS; and regions outside the CDS span of any gene model. Regions in the last class may still lie in untranslated regions or in non-coding gene models. Each candidate region was graded by its evidence. High-evidence regions were supported at the gathering threshold by miniprot alignments of at least two different queries from at least two species. Other regions supported at the gathering threshold by miniprot alignments were graded as medium. Low-evidence regions were supported only at the E-value threshold or only by six-frame translation. Candidate regions were reported separately from the gene counts. As a positive control for sensitivity, the search was repeated for each genome with the queries of that species removed, and the proportion of annotated Kunitz/BPTI-positive genes recovered at the gathering threshold was recorded.

### Second public annotations of the mosquito and tick assemblies

Ensembl Metazoa release 63 [18] provides gene sets for the same assemblies of *A. aegypti* (AaegL5.3) and *I. scapularis* that were used in RefSeq. GFF3 files and protein sets were downloaded and checked against the published checksums. Sequence names were converted to RefSeq accessions with the NCBI assembly reports. Before the family comparison, the overlap of the two gene sets was measured. The first measure was the proportion of Ensembl genes whose CDS span was identical to that of at least one RefSeq gene. The second was the proportion of Ensembl genes whose set of CDS segment coordinates, pooled across transcripts, was identical to that of at least one RefSeq gene. Strand and one-to-one correspondence were ignored. Family counts and gene correspondence were then determined as described above for the leech.

## Results

### Protein entries and genes in eight genomes

Seven protease-inhibitor families were counted in the protein sets of eight animal genomes. A total of 1,296 protein entries carried at least one of the seven domains. Because some entries carried more than one domain type, they gave 1,346 entry–family assignments in 48 species–family combinations with at least one positive gene (Fig. 1a; Additional file 2). In 36 of the 48 combinations, the number of protein entries was larger than the number of genes. The largest difference was found for Kunitz/BPTI in *A. aegypti*, in which 23 protein entries belonged to 5 genes (4.60 entries per gene). For Kunitz/BPTI, the ratio was 2.61 in *D. rotundus*, 1.79 in *C. elegans*, 1.66 in *N. americanus*, 1.54 in *D. melanogaster*, 1.25 in *H. manillensis*, 1.03 in *I. scapularis* and 1.00 in *C. teleta* (Fig. 1b). These results indicate that the effect of the counting unit differs between species and between families of the same species.

**Fig. 1.**
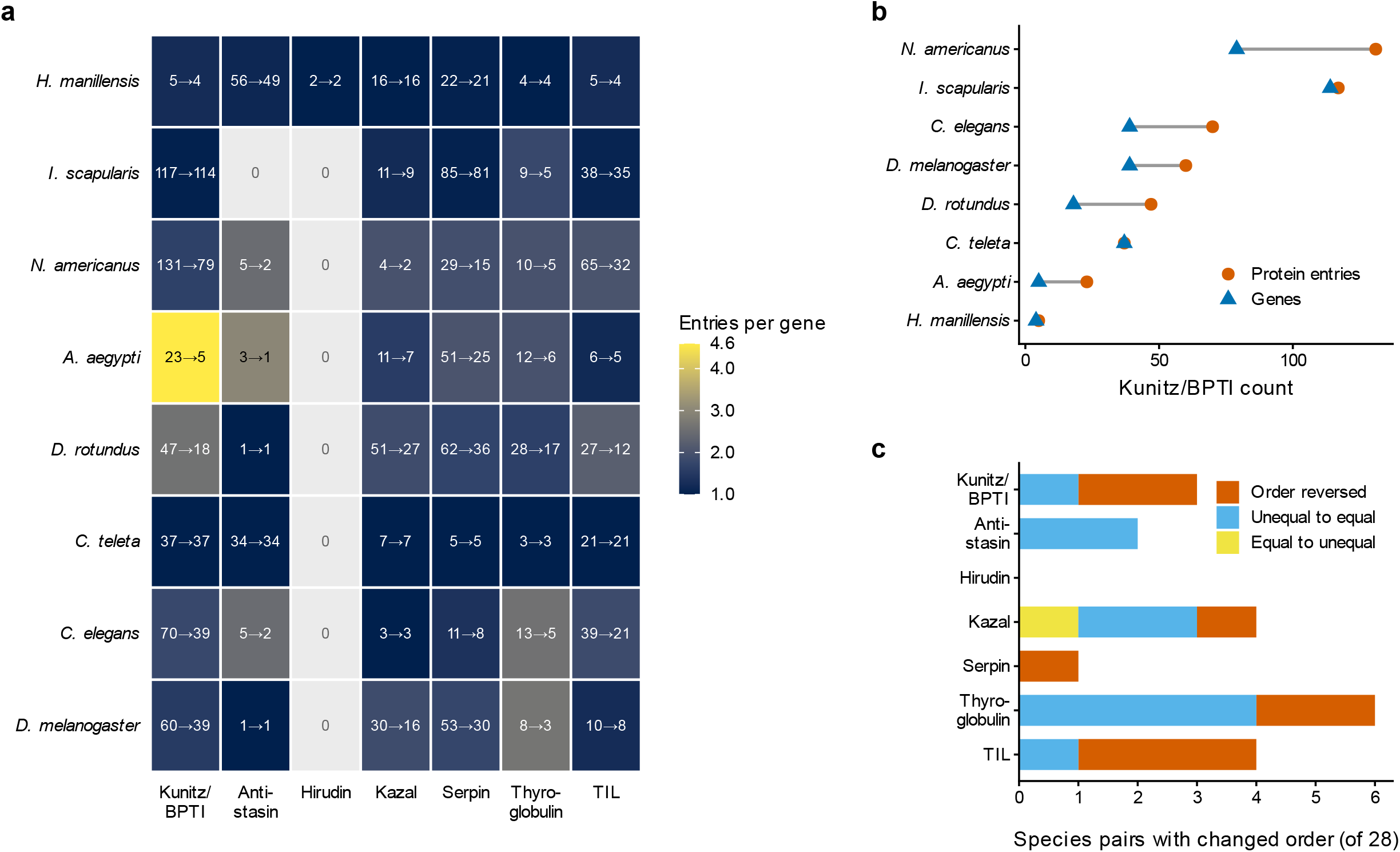
Protein-entry and gene counts of seven protease-inhibitor families in eight genomes. **a** Number of domain-positive protein entries and genes for each family and species (entries→genes). Colour shows the number of entries per gene; grey cells contain no positive gene. **b** Kunitz/BPTI counts per species. Circles, protein entries; triangles, genes (longest entry per gene). **c** Number of species pairs (of 28 per family) whose order changed between protein entries and genes: order reversed, two unequal counts became equal, or two equal counts became unequal. Genes in **a**–**c** were counted as positive when their longest entry carried the domain. Domains were detected with Pfam profile HMMs at the gathering thresholds.

Interestingly, the change of counting unit also changed the order of species. For each family, the 28 species pairs were compared under the two counting units. Of the 196 comparisons, the order was reversed in 9, two unequal counts became equal in 10, and two equal counts became unequal in 1 (Fig. 1c). For Kunitz/BPTI, *N. americanus* had more protein entries than *I. scapularis* (131 vs 117) but fewer genes (79 vs 114). Similarly, *D. rotundus* had more protein entries than *C. teleta* (47 vs 37) but fewer genes (18 vs 37). Order reversals were also found for TIL (3 pairs), Thyroglobulin (2), Kazal (1) and Serpin (1).

Gene counts were also compared under two rules for assigning domain positivity to a gene. A gene was scored as positive either when its longest entry carried the domain (used above) or when any of its entries carried it. Under the second rule, the count increased in only four species–family combinations: Antistasin in *H. manillensis* (49 to 50), Kunitz/BPTI in *N. americanus* (79 to 82), TIL in *N. americanus* (32 to 34) and Kazal in *D. rotundus* (27 to 28). None of these changes altered the order of any species pair. Thus, the rule used to assign domain positivity to genes had little effect on the counts or on the order of species.

### Annotated transcripts and neighbouring duplicate genes

To test whether the protein entries of one gene belonged to a single annotated locus, the CDS coordinates of each entry were compared with those of the representative entry. In total, 213 genes had two or more domain-positive entries in at least one family (223 gene–family records). In 218 of the 223 records (208 genes), all entries shared CDS coordinates with the representative entry. The remaining five records comprised three genes of *N. americanus* and one gene each of *D. rotundus* and *C. elegans* (Additional file 3). In each of them, at least one entry had no CDS in common with the representative entry, but all entries lay within the annotated gene feature. In addition, 107 of the 1,346 entry–family assignments were entries with an amino acid sequence identical to another entry of the same gene. For example, 11 of the 23 Kunitz/BPTI entries of *A. aegypti* were identical copies of other entries.

Neighbouring genes with nearly identical proteins were counted separately. Under the main criterion (at least 98% identity, at most 20 kb apart, no overlap), nine candidate tandem duplicate pairs were identified (Additional file 3). These included two Kunitz/BPTI pairs and two Serpin pairs in *I. scapularis*, one Kunitz/BPTI pair and one Serpin pair in *N. americanus*, and three Antistasin pairs in *C. teleta*. Under the permissive criterion, 34 pairs were found. No candidate pair was found in *A. aegypti*, so its 4.6-fold difference came from several entries of the same annotated genes. In *H. manillensis*, two pairs of overlapping gene models with similar proteins were found among the Antistasin genes. Both pairs consisted of one model from each track of the published annotation (see below).

### Two annotations of the same leech assembly

The two annotations of the *H. manillensis* assembly contained 22,836 and 20,377 gene models, and their domain-positive genes were compared by genomic position (Fig. 2a; Additional file 4). The GALBA annotation contained 3 Kunitz/BPTI-, 11 Antistasin-, 12 Kazal-, 18 Serpin-, 5 Thyroglobulin- and 4 TIL-positive genes and no Hirudin-positive gene. For Kunitz/BPTI, Serpin, TIL and Thyroglobulin, most positive genes of the published annotation had a one-to-one partner in the GALBA annotation. The proportions were 3 of 4 for Kunitz/BPTI, 17 of 21 for Serpin, 4 of 4 for TIL and 4 of 4 for Thyroglobulin, and the remaining Kunitz/BPTI gene had no same-strand GALBA gene model.

**Fig. 2.**
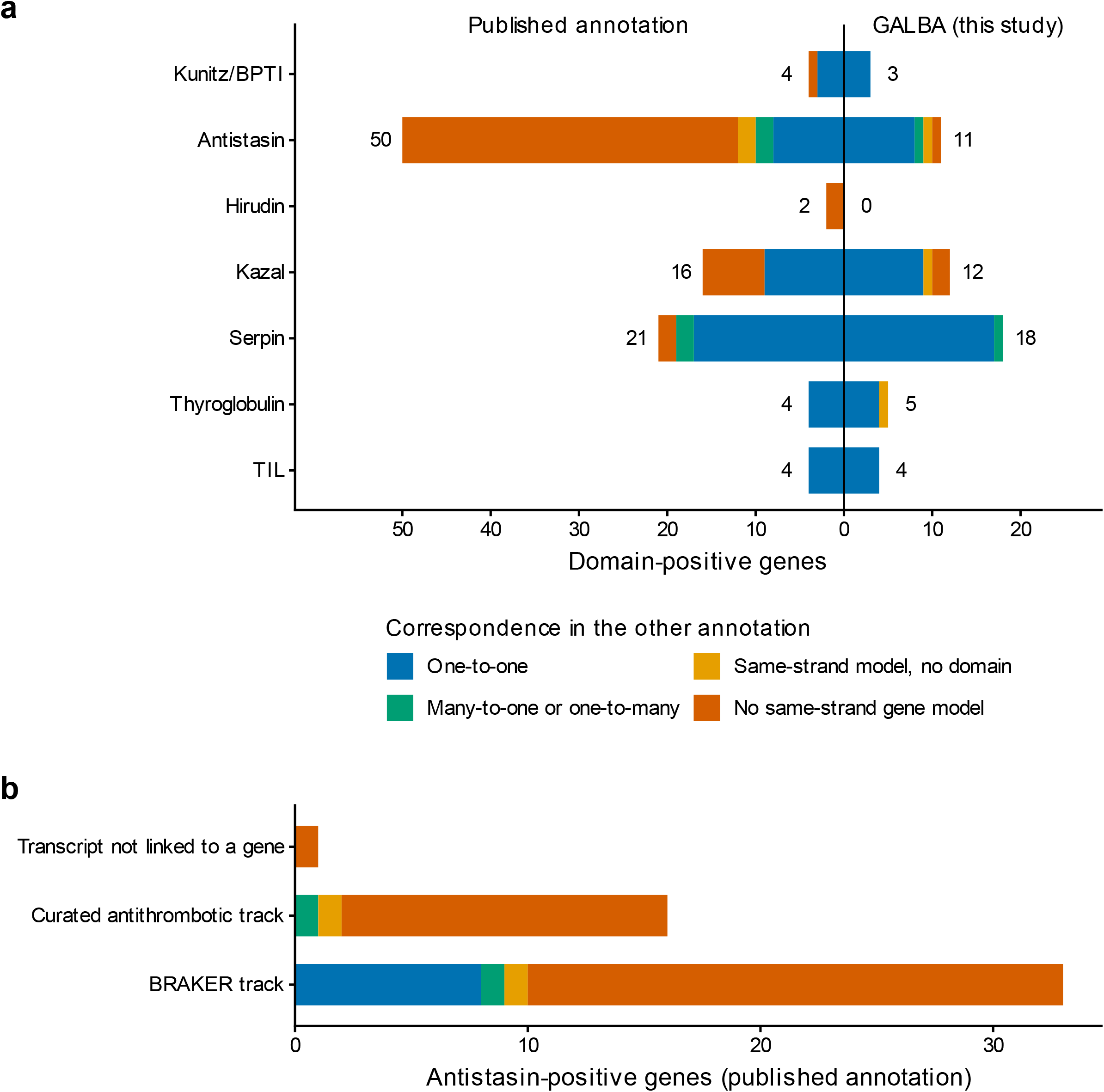
Domain-positive genes in two annotations of the same *H. manillensis* assembly. **a** Positive genes of each family in the published annotation (left) and the GALBA annotation generated in this study (right), classified by their correspondence in the other annotation. Numbers show the total positive genes on each side. **b** Antistasin-positive genes of the published annotation by annotation track. Colours as in **a**. Genes were scored as positive when any entry carried the domain and were compared by the overlap of their CDS spans on the same strand. In **b**, the curated transcript model without a linked gene feature is shown separately.

In contrast, larger differences were found for Antistasin and Kazal. Of the 50 Antistasin-positive genes of the published annotation, 8 had a one-to-one partner and 2 formed a many-to-one overlap with one GALBA positive gene. Another 2 overlapped a GALBA model without the domain on the same strand, and the remaining 38 had no GALBA gene model on the same strand. Of the 16 Kazal-positive genes, 7 had no same-strand GALBA gene model. Notably, both Hirudin-positive genes of the published annotation had no same-strand GALBA gene model.

The GALBA annotation also contained positive genes at positions where the published annotation had no same-strand gene model or a model without the domain (Fig. 2a). One Antistasin-positive gene and two Kazal-positive genes lay at positions without a published gene model on the same strand. One Antistasin, one Kazal and one Thyroglobulin gene lay at positions where the published model lacked the domain.

The published annotation combines a BRAKER track with a track of curated antithrombotic genes. Of the 16 Antistasin-positive gene models of the curated track, 14 had no same-strand GALBA gene model, compared with 23 of the 33 positive genes of the BRAKER track (Fig. 2b). One further Antistasin-positive transcript model of the curated track also had no same-strand GALBA gene model. This model lacks a linked gene feature in the published GFF3 file. Four curated Antistasin genes overlapped a positive gene of the BRAKER track, and in two of these cases the proteins were 91% and 100% identical. Thus, some Antistasin loci are represented by two overlapping gene models in the published annotation.

### Kunitz/BPTI candidate regions outside the annotations

To test whether the annotations missed Kunitz/BPTI genes, the eight genome assemblies were searched without using the gene models. When the complete query set was used, which included the proteins of each target species, a Kunitz/BPTI domain was detected on the same-strand CDS of every annotated Kunitz/BPTI-positive gene (338 of 338 genes). When the queries of the target species were removed, the proportion of annotated genes recovered ranged from 31% in *I. scapularis* (35 of 114) and 36% in *D. melanogaster* (14 of 39) to 83% in *D. rotundus* (15 of 18) (Fig. 3a; Additional file 5). These results indicate that queries from other species detect only part of a divergent family.

**Fig. 3.**
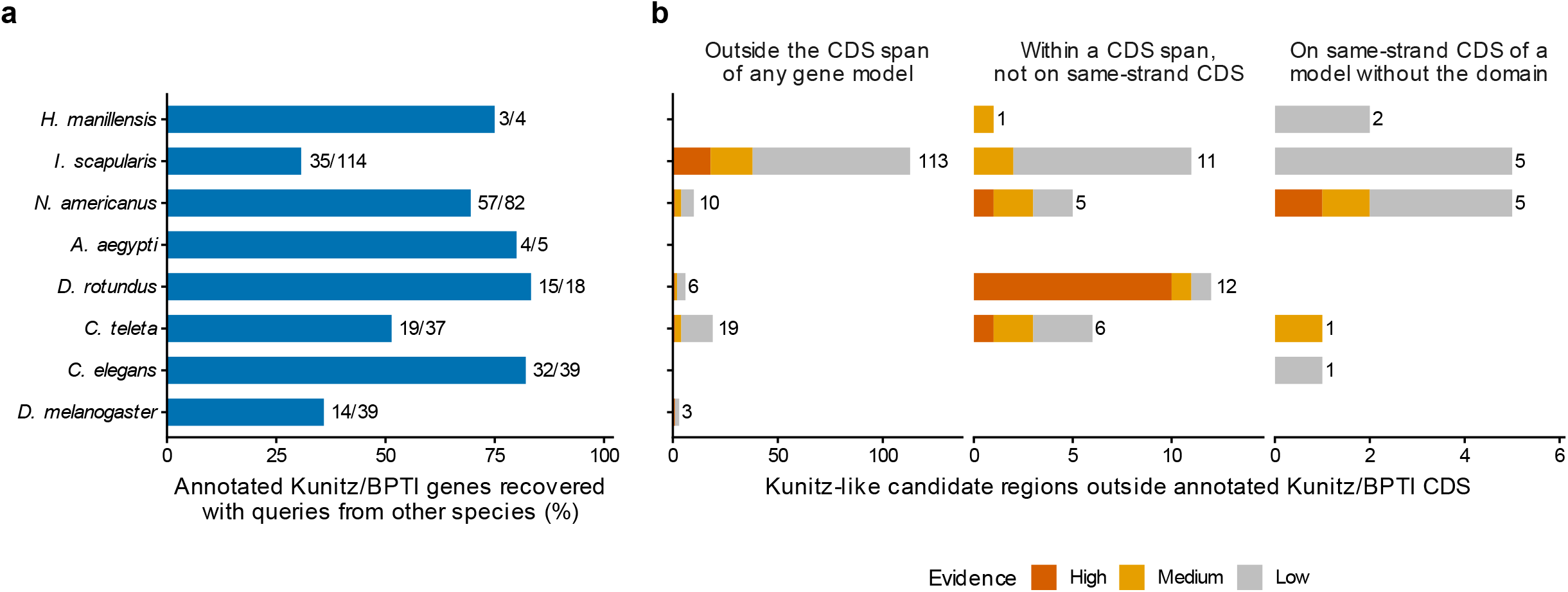
Search of the eight genome assemblies for Kunitz/BPTI domains without using gene models. **a** Proportion of annotated Kunitz/BPTI-positive genes (genes with any positive entry) detected when the queries of the target species were removed from the query set (gathering threshold). Numbers give recovered and annotated genes. With the complete query set, which included the proteins of each species, all 338 annotated genes were detected. **b** Kunitz-like candidate regions outside the same-strand CDS of annotated Kunitz/BPTI-positive genes, classified by position: outside the CDS span of any gene model; within the CDS span of a gene model (either strand) but not on its same-strand CDS; and on the same-strand CDS of a gene model without the domain. Bars include low-evidence regions, and each panel has its own scale. Numbers give the total number of regions. Evidence: high, supported at the gathering threshold by miniprot alignments of at least two different queries from at least two species; medium, other regions supported at the gathering threshold by miniprot alignments; low, supported only at the E-value threshold or only by six-frame translation. Candidate regions were reported separately from the gene counts.

Kunitz-like regions outside the CDS of annotated Kunitz/BPTI genes were classified by their position relative to the gene models (Fig. 3b). Regions with high or medium evidence outside the CDS span of any gene model were found mainly in *I. scapularis* (38 regions; 18 high, 20 medium). Fewer such regions were found in *N. americanus* (4), *C. teleta* (4), *D. rotundus* (2) and *D. melanogaster* (1). Regions with high or medium evidence within the CDS span of a gene model but not on its same-strand CDS were found in *D. rotundus* (11), *N. americanus* (3), *C. teleta* (3), *I. scapularis* (2) and *H. manillensis* (1). In *D. rotundus*, nine of these regions were inside one Kunitz/BPTI-positive gene model (GeneID 128780253). This model spans 316 kb and has 13 annotated protein entries, all of which carry the domain. Few regions overlapped the CDS of a gene model without the domain (two in *N. americanus* and one in *C. teleta* with high or medium evidence). No candidate region was found in *A. aegypti*, and only low-evidence regions were found in *C. elegans*.

### Second public annotations of the mosquito and tick assemblies

Ensembl Metazoa provides a second gene set for the same assemblies of *A. aegypti* and *I. scapularis*. Before family counts were compared, the two gene sets of each species were compared as a whole. In *A. aegypti*, 98.9% of the Ensembl genes had a CDS span identical to that of a RefSeq gene, and 97.2% had identical coordinates for all CDS segments. In *I. scapularis*, both values were 99.8%. Thus, in these two species the coordinates of the two public gene sets overlap almost completely. Consistent with this, the Kunitz/BPTI counts were identical in the two annotations of each species (*A. aegypti*, 23 protein entries and 5 genes; *I. scapularis*, 117 entries and 114 genes; Additional file 6). Among the other families, the two annotations of *A. aegypti* differed by one gene for Kazal and TIL, and all other counts were identical. The comparison of annotation sources was therefore based on the two leech annotations.

## Discussion

Counts of gene family members are widely used to compare genomes, and several recent studies have annotated anticoagulant or antithrombotic genes in leech genomes [1, 2, 10, 19, 20]. Such counts depend on how protein entries are assigned to genes and on how the genes were annotated. In this study, we thus measured both effects for seven protease-inhibitor families in eight animal genomes.

The counting unit changed the counts by different factors in different species and families. Collapsing protein entries to genes reduced the Kunitz/BPTI count 4.6-fold in *A. aegypti* but left it unchanged in *C. teleta*, and the order of two species was reversed in 9 of 196 pairwise comparisons. The rule used to assign domain positivity to a gene had little effect on these differences. One transcript per gene is commonly used for orthology inference [3], and our results show that the same step is needed for simple domain counts. Without it, the comparison partly reflects how many transcripts each annotation project modelled. In addition, candidate tandem duplicates with nearly identical proteins were rare under our criteria, and in 218 of 223 gene–family records all entries shared CDS coordinates with the representative entry. Most extra entries are therefore transcript models of the same annotated locus. However, transcript evidence is needed to show whether they are real alternative transcripts.

The annotation source affected the counts in a family-specific way. For Kunitz/BPTI, Serpin, TIL and Thyroglobulin, most positive genes of one leech annotation had a one-to-one partner in the other. In contrast, 38 of the 50 Antistasin-positive genes and both Hirudin-positive genes of the published annotation had no same-strand gene model in the GALBA annotation. The proportion without a same-strand GALBA model was higher for the curated antithrombotic track (15 of 17 models, including one transcript model without a linked gene feature) than for the BRAKER track (23 of 33) of the published annotation [2]. Short, secreted and lineage-specific proteins such as hirudin may be difficult to predict from distant protein evidence alone, and family-targeted curation can add such genes. However, the curated track also contained gene models that overlapped BRAKER models at the same position. The GALBA annotation also contained Antistasin and Kazal genes at positions where the published annotation had no same-strand model or a model without the domain. Thus, counts of these families from different annotations are not directly comparable. The antithrombotic gene counts reported in different leech genome studies [2, 10] should therefore be read together with the annotation procedure of each study.

Because the genome search recovered every annotated Kunitz/BPTI-positive gene when proteins of the same species were included, the candidate regions mark positions where the annotations may have missed members of the family. Such regions were most numerous in *I. scapularis*, in which the Kunitz/BPTI family has expanded [8]. In *D. rotundus*, most of them lay within the CDS span of one long gene model.

A comparison of annotation sources is informative only when the annotations contain different gene models. For the mosquito and the tick, the second public gene sets shared nearly all gene-model coordinates with RefSeq and gave identical Kunitz/BPTI counts. Annotations distributed by different databases may therefore share most of their gene models, and this overlap should be checked before a comparison of annotation sources is interpreted.

This study has several limitations. Domain detection used one profile HMM per family with the Pfam gathering thresholds, and members with highly divergent or partial domains may be missed. The genome search covered only Kunitz/BPTI. Its candidate regions may be missed genes, pseudogenes, gene fragments or additional exons of existing genes, and their structure and expression remain to be tested. Because queries from other species recovered only part of the annotated genes, the absence of candidates in a genome does not show that its annotation is complete, especially for rapidly diverging families such as the tick Kunitz/BPTI family [8]. Gene models and their correspondence between annotations were analysed as annotated positions without experimental validation, and the secretion and inhibitory activity of the domain-positive proteins remain to be examined. Only one assembly, that of *H. manillensis*, had two annotations with substantially different gene models. The eight genomes were chosen because their annotations come from different sources, and they are not a sample for testing whether any trait, such as blood feeding, is associated with family size. Distinguishing orthologues from paralogues requires gene-tree or gene-family models [3], which were outside the scope of this study. However, these results illustrate sources of variation that can affect comparisons based on annotated protein sets.

Based on these results, we suggest four practices for comparing gene family sizes across species. First, each gene should be counted once, and the rule used to assign domain positivity to genes (for example, any entry or the longest entry) should be reported. Second, the source and version of each annotation should be reported. Third, the annotations should be checked with a search of the genome assembly, and the sensitivity of the search should be reported. Fourth, the overlap of gene models should be checked before two annotations are used to compare annotation pipelines.

## Conclusions

Family counts taken from public protein sets changed with the counting unit and with the annotation source. Collapsing protein entries to genes changed the counts of seven protease-inhibitor families by up to 4.6-fold and reversed the order of species in several comparisons. Two annotations of the same leech assembly gave different sets of Antistasin-, Kazal- and Hirudin-positive genes. These results provide useful information for comparing gene family sizes between genomes annotated by different projects.

## Abbreviations

BPTI: bovine pancreatic trypsin inhibitor
BUSCO: Benchmarking Universal Single-Copy Orthologs
CDS: coding sequence
HMM: hidden Markov model
TIL: trypsin inhibitor-like

## Declarations

### Ethics approval and consent to participate

Not applicable. This study used only public genome data and published annotations.

### Consent for publication

Not applicable.

### Availability of data and materials

All genome assemblies and annotations are publicly available from NCBI under the accessions listed in Table 1 and from Ensembl Metazoa release 63 [18]. The leech coding sequences and annotation were released by Liu et al. [2] (figshare doi:10.6084/m9.figshare.24187377). The result tables generated in this study are provided in Additional files 1–6. The data and R scripts used to draw Figs. 1–3 are available at Zenodo (doi:10.5281/zenodo.23757292). The GALBA annotation of the *H. manillensis* assembly and the analysis scripts are available from the corresponding author on request.

### Authors’ contributions

Fang Zhao and Feng Zhao conceived the study. Fang Zhao and Juan Zhao curated the data. Fang Zhao, Juan Zhao and Feng Zhao performed the formal analysis. Fang Zhao prepared the figures. Fang Zhao and Feng Zhao wrote the original draft, and all three authors reviewed and edited the manuscript. Feng Zhao acquired the funding and supervised the study. All authors read and approved the final manuscript.

## Acknowledgements

Not applicable.

## Additional files

**Additional file 1.** File name: Additional_file_1.xlsx; format: Microsoft Excel. Title: Settings of the GALBA annotation of the *H. manillensis* assembly. Description: protein evidence, run settings and training correction, and the correspondence of sequence names between the GenBank assembly and the published annotation.

**Additional file 2.** File name: Additional_file_2.xlsx; format: Microsoft Excel. Title: Family counts under three counting units. Description: counts of seven protease-inhibitor families as protein entries, genes with any positive entry and genes with a positive longest entry in eight genomes, and species pairs whose order changed between protein entries and genes.

**Additional file 3.** File name: Additional_file_3.xlsx; format: Microsoft Excel. Title: Genes with several positive protein entries and candidate tandem duplicates. Description: CDS relations between the entries of each gene and the representative entry, and neighbouring gene pairs under the main and permissive criteria.

**Additional file 4.** File name: Additional_file_4.xlsx; format: Microsoft Excel. Title: Correspondence of positive genes between two leech annotations. Description: classification of each domain-positive gene of the published and the GALBA annotations of the *H. manillensis* assembly, with CDS coordinates and overlapping genes.

**Additional file 5.** File name: Additional_file_5.xlsx; format: Microsoft Excel. Title: Genome search for Kunitz/BPTI domains. Description: summary by species, positive control and all candidate regions with their domain segments, class and evidence.

**Additional file 6.** File name: Additional_file_6.xlsx; format: Microsoft Excel. Title: RefSeq and Ensembl Metazoa gene sets of *A. aegypti* and *I. scapularis*. Description: overlap of gene-model coordinates, family counts and correspondence of positive genes.

